# Theta-gamma coupling depends on breathing rate

**DOI:** 10.1101/2020.10.22.349936

**Authors:** Maximilian Hammer, Chrysovalandis Schwale, Jurij Brankačk, Andreas Draguhn, Adriano BL Tort

## Abstract

Temporal coupling between theta and gamma oscillations is a hallmark activity pattern of several cortical networks and becomes especially prominent during REM sleep. In a parallel approach, nasal breathing has been recently shown to generate phase-entrained network oscillations which also modulate gamma. Both slow rhythms (theta and respiration-entrained oscillations) have been suggested to aid large-scale integration but they differ in frequency, display low coherence, and modulate different gamma sub-bands. Respiration and theta are therefore believed to be largely independent. In the present work, however, we report an unexpected but robust relation between theta-gamma coupling and respiration in mice. Interestingly, this relation takes place not through the phase of individual respiration cycles, but through respiration rate: the strength of theta-gamma coupling exhibits an inverted V-shaped dependence on breathing rate, leading to maximal coupling at breathing frequencies of 4-6 Hz. Noteworthy, when subdividing sleep epochs into phasic and tonic REM patterns, we find that breathing differentially relates to theta-gamma coupling in each state, providing new evidence for their physiological distinctiveness. Altogether, our results reveal that breathing correlates with brain activity not only through phase-entrainment but also through rate-dependent relations with theta-gamma coupling. Thus, the link between respiration and other patterns of cortical network activity is more complex than previously assumed.

## Introduction

The coupling between theta phase (4-12 Hz) and the amplitude of gamma (30-160 Hz) has been perhaps the most studied case of cross-frequency coupling between neuronal network oscillations to date (Lisman and Jensen, 2013). This phase-amplitude coupling is thought to serve as a means of integrating locally generated gamma assemblies across distant regions (Canolty and Knight, 2010); that is, while gamma is believed to be a physiological marker of local computations (Sirota et al., 2008; Buzsáki and Wang, 2012), theta tends to be coherent across multiple structures and is thus considered a global brain signal well suited for linking distributed gamma activity (Tort et al., 2007; Colgin, 2011; Hyafil et al., 2015). In support of this possibility, theta-gamma coupling has been observed in several subcortical and neocortical regions (Chrobak and Buzsáki, 1998; Tort et al., 2008, 2009; Scheffzük et al., 2011; Stujenske et al., 2014).

The study of cross-frequency coupling revealed that gamma oscillations, once thought as a monolithic entity, may be divided into specific sub-bands that correspond to different physiological processes (Belluscio et al., 2012; Lasztóczi and Klausberger, 2014; Schomburg et al., 2014; Bott et al., 2016; Scheffer-Teixeira and Tort, 2017). Importantly, the coupling between theta and the gamma sub-bands is not a static phenomenon but rather depends on the recorded region, cognitive demands and behaviors (Tort et al., 2008; Scheffer-Teixeira et al., 2012; Schomburg et al., 2014; Bott et al., 2016; Bandarabadi et al., 2019), and is likely to mark functional brain states (Lopes-Dos-Santos et al., 2018; Zhang et al., 2019). In particular, in the neocortex of mice and rats, theta-gamma coupling is most prominent during REM sleep (Scheffzük et al., 2011; Brankačk et al., 2012; Scheffer-Teixeira and Tort, 2017), where it is believed to underlie the processing of information acquired during awake states (Bandarabadi et al., 2019; see also Boyce et al., 2016).

A more recent line of investigation has shown that nasal breathing induces local field potential (LFP) oscillations at the same frequency as respiration in several brain regions not primarily related to olfaction (for reviews, see Tort et al., 2018a; Heck et al., 2019). Respiration phase is also capable of modulating the amplitude of gamma-frequency oscillations (Ito et al., 2014; Yanovsky et al., 2014; Rojas-Líbano et al., 2018; Cavelli et al., 2020), especially in frontal regions (Biskamp et al., 2017; Zhong et al., 2017). Interestingly, the gamma sub-band modulated by the respiration phase differs from the gamma sub-bands modulated by the theta phase, suggesting that different slow rhythms use specific frequency ranges of fast oscillations for their local influence (Zhong et al., 2017). The so-called respiration-entrained oscillations could thus serve as a substrate for large scale integration of information across the brain (Heck et al., 2017; Tort et al., 2018a; Heck et al., 2019), in similarity to what has been previously proposed for other slow network rhythms such as theta (Isomura et al., 2006; Canolty and Knight, 2010).

Previous findings have strongly suggested that theta and respiration rhythms are independent of one another (Lockmann et al., 2016; Nguyen Chi et al., 2016). This is because, in addition to the gamma sub-band selectivity mentioned above (Zhong et al., 2017), theta and respiration often have different frequencies and are not phase coherent, and, moreover, the rhythms have clearly distinct mechanisms of generation and regional specificities (Yanovsky et al., 2014; Lockmann et al., 2016; Nguyen Chi et al., 2016; Tort et al., 2018b). However, it should be noted that these studies have mainly focused on entrainment-relations involving the *respiration phase*. Here, we took a new approach and investigated whether the *respiration rate* – not phase – would relate theta and gamma activity. The rationale behind this approach is that changes in respiration rate could potentially reflect changes in internal brain states, as inferred by network oscillatory patterns. To control for gross behavioral variables as confounding factors (i.e., locomotor activity), we have focused on immobile animals. In specific, we analyzed respiration and LFPs from the posterior parietal cortex of mice recorded during REM sleep in a whole-body plethysmograph (Figure 1A,B). In this behavioral state, the posterior parietal cortex exhibits prominent theta and gamma oscillations and theta-gamma coupling (Figure 1C-F; see also Scheffzük et al., 2011; Brankačk et al., 2012; Scheffer-Teixeira and Tort, 2017). Our aim was therefore to investigate if the instantaneous breathing rate influences this hallmark oscillatory network pattern.

**Figure 1.**
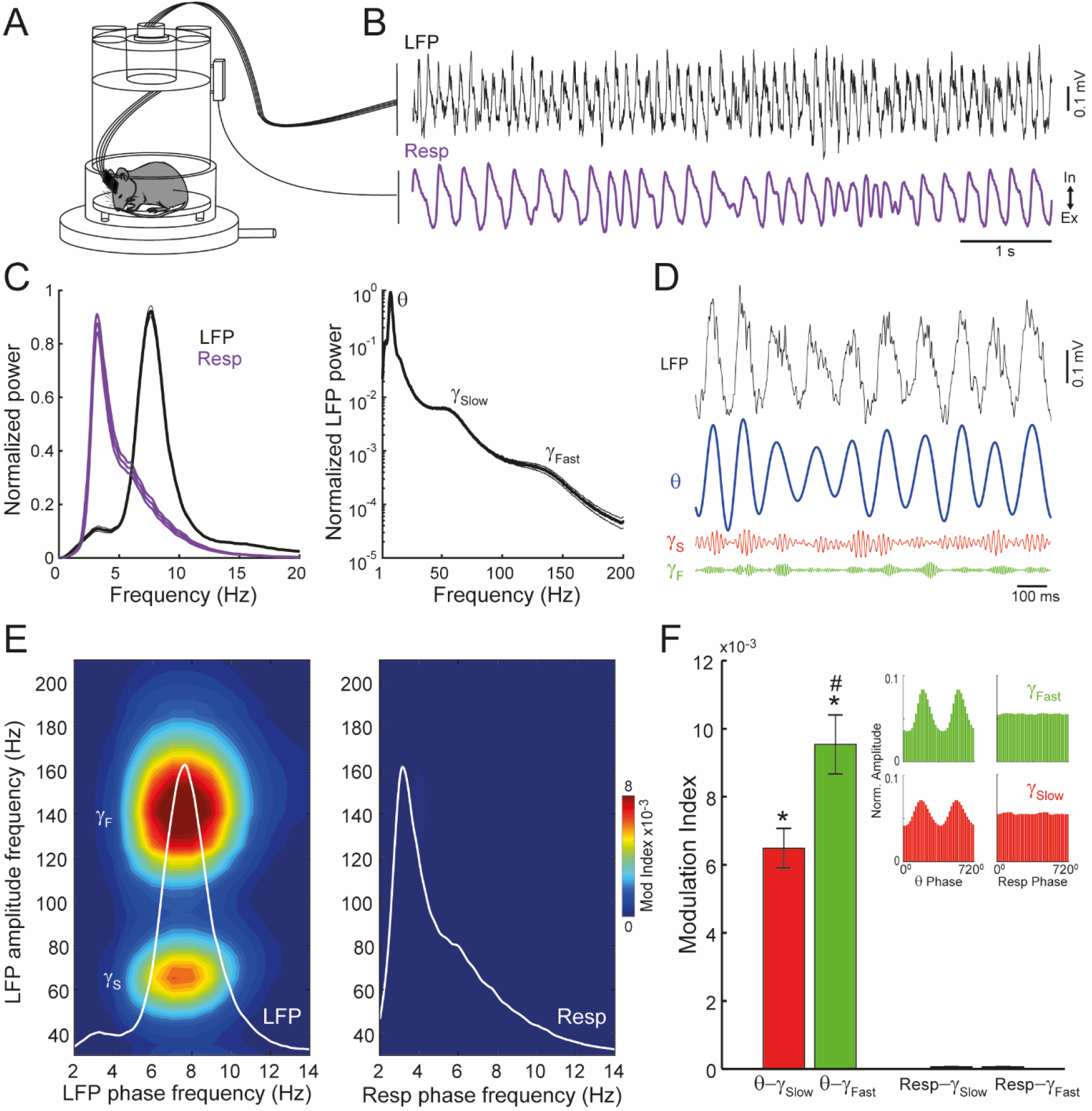
During REM sleep, theta, but not respiration, modulates fast and slow gamma activity. (A) Schematic depiction of a mouse recorded inside the plethysmograph during sleep. (B) Example of parietal local field potential (LFP) and respiration (Resp) recordings during REM sleep. Notice prominent theta (θ) oscillations and variable breathing rate. (C) (Left) Normalized LFP and Resp power spectra (mean ± SEM; n=22 mice). The normalization was obtained by dividing the spectrum of each signal by the maximal power value. (Right) Normalized LFP power spectrum in logarithm scale, revealing power bumps at the slow (γS) and fast (γF) gamma sub-bands. (D) Example of raw and filtered LFP traces (θ: 4-12 Hz; γS: 50-80 Hz; γF: 120-160 Hz). Notice rhythmical variations of gamma amplitude. (E) Average REM sleep comodulogram (n=22 mice) computed for the amplitude of parietal LFP oscillations and the phase of either the parietal LFP (left) or Resp (right). The overlaid white lines depict the corresponding power spectra. The LFP theta phase modulates both gamma sub-bands, while the phase of breathing cycles has no direct influence over parietal gamma activity during sleep. (F) Mean modulation index for the amplitude of the two gamma sub-bands and the phase of theta or respiration, as labeled. The inset plots show example phase-amplitude distributions for a representative animal. ^#^p=0.0016, paired t-test between θ-γS and θ-γF; *p<10^−8^ paired t-tests between θ-γS and Resp-γS and between θ-γF and Resp-γF.

## Results

### Prominent neocortical theta-gamma coupling during REM sleep

We started by first characterizing the patterns of oscillatory activity in the posterior parietal cortex of mice during REM sleep within the whole-body plethysmograph (Figure 1A,B). On average, theta frequency was faster than the breathing rate derived from the plethysmograph signal (7.6 Hz vs 3.2 Hz; Figure 1C left). In addition to the expected, large-amplitude theta rhythm, REM sleep was also characterized by parietal LFP activity in two gamma sub-bands, as inferred by a power bump in the LFP spectrum ranging from ~50 to ~80 Hz and another from ~120 to ~160 Hz (Figure 1C right). In this work, we refer to these fast LFP activities as slow- and fast-gamma oscillations, respectively. Consistent with previous reports in mice (Scheffzük et al., 2011, 2013; Brankačk et al., 2012), during REM sleep the instantaneous amplitude of both gamma sub-bands was strongly modulated by theta phase (Figure 1D,E). Theta-fast gamma coupling was statistically significantly higher than theta-slow gamma coupling (t(21)=3.62, p=0.0016, paired t-test; Figure 1F). Of note, although respiration phase has also been shown to modulate gamma amplitude (Ito et al., 2014; Yanovsky et al., 2014), this effect is most prominent in frontal regions during quiet immobility (Biskamp et al., 2017; Zhong et al., 2017). In the current dataset, no meaningful coupling to respiration phase occurred for gamma amplitude in the posterior parietal cortex during REM sleep (Figure 1E,F).

### Theta-gamma coupling depends on breathing rate

We next sought to investigate whether respiration could influence brain activity in different ways to the already described phase-entrainment relations (Tort et al., 2018a). To that end, we tracked the breathing rate during REM sleep by computing normalized power spectra for the plethysmograph-derived respiration signal using non-overlapping windows of 500 ms (Figure 2A). The normalization was performed only to aid visualization and was obtained by dividing each power spectrum by its maximal value, so that, for each 500-ms window, the maximal power value is 1; the frequency of maximal respiration power was then taken as the breathing rate of the window (Figure 2A left). During REM sleep, the average mode of the individual breathing rate distributions was 3.33 ± 0.20 Hz (M ± SD, n = 22 animals; pooled distribution mode: 3.35 Hz), while the average breathing rate was 4.27 ± 0.38 Hz (pooled distribution average: 4.35 Hz). The mean was higher than the mode due to the skewed distribution of breathing rates towards faster frequencies (Figure 2A right). In fact, breathing rates up to 11 Hz could also be observed, though the 25th and 75th quartiles of the pooled distribution were 3.10 and 5.05 Hz, respectively (Figure 2A right); the mean (±SD) interquartile range of individual distributions was 1.86 ± 0.49 Hz.

**Figure 2.**
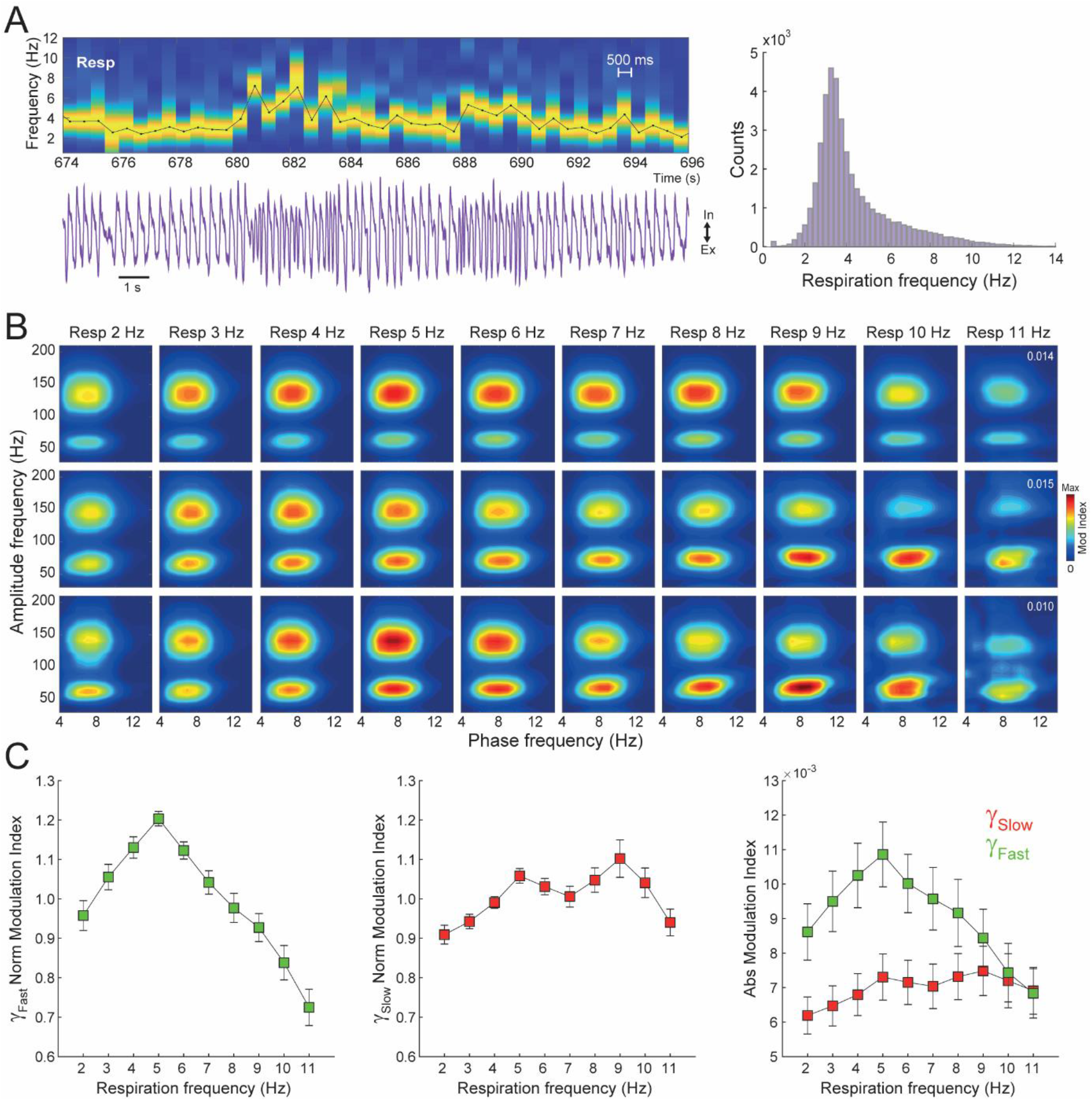
Theta-gamma coupling depends on breathing rate. (A) (Left) Example time-frequency power spectrogram for the Resp signal using non-overlapping 500-ms windows, which was used to track breathing rate. The power spectrum of each window was normalized by the maximal value. The black line depicts the breathing rate, as defined by the peak power frequency. (Right) Histogram counts of respiration peak frequency in 500-ms windows (pool over 22 animals). (B) Phase-amplitude comodulograms computed for REM sleep LFP epochs binned by respiration frequency. The title of each column states the average breathing rate (respiration frequency intervals: 1-3 Hz, 2-4 Hz, 3-5 Hz, 4-6 Hz, 5-7 Hz, 6-8 Hz, 7-9 Hz, 8-10 Hz, 9-11 Hz, and >10 Hz). Each row shows the results of a different animal. The inset numbers in the rightmost comodulograms indicate the maximum of the modulation index color scale. Notice similar patterns among animals and that both theta-fast gamma and theta-slow gamma coupling depend on breathing rate. (C) Group data of normalized (left and middle panels) and absolute (right panel) theta-gamma coupling strength as a function of respiration frequency (mean ± SEM; n=22 mice). The normalization was performed within animals by dividing the modulation index values by its mean. See text for statistical analyses.

We next proceeded to evaluate whether the prominent theta-gamma coupling observed in the posterior parietal cortex during REM sleep (Figure 1E,F) would depend on respiration rate. To that end, we computed phase-amplitude comodulation maps using LFP epochs binned into fixed respiration frequency intervals (see Materials and Methods for details). Unexpectedly, the results revealed an inverted V-shaped relation between the strength of theta-fast gamma coupling and breathing rate during sleep, in which coupling strength was maximal when animals breathed around 5 Hz, and linearly decreased from there for either increases or decreases of breathing rate relative to 5 Hz (Figure 2B,C). A repeated measures ANOVA confirmed a highly significant effect of respiration rate on theta-fast gamma coupling strength (F(9,203)=17.41, p<10^−20^). Interestingly, theta-slow gamma coupling strength also depended on breathing rate but exhibited a different profile, namely, an asymmetrical, inverted W-shaped relation, with peak coupling when animals breathed either around 5 Hz or 9 Hz (Figure 2B,C). The effect of respiration on theta-slow gamma coupling was highly significant (F(9,203)=4.55, p<0.0001, repeated measures ANOVA), and so was the profile difference in breathing rate dependence between slow- and fast-gamma coupling to theta (F(9,406)=10.39, p<10^−50^, two-way ANOVA, interaction effect). In all, these findings show that, in addition to the well-known phase-locking effects (Tort et al., 2018a), breathing also relates to oscillatory brain activity through its instantaneous rate.

### Breathing rate differentially relates to theta-gamma coupling in phasic and tonic REM sleep

The microstructure of REM sleep reveals a heterogeneous state with regard to neurophysiological features such as eye movement, ponto-geniculo-occipital spikes, muscle twitches, and vegetative arousal (Sakai et al., 1973; Blumberg et al., 2020; Simor et al., 2020). This heterogeneity motivated the division of REM sleep into tonic and phasic states. In rodents, the latter is characterized by phasic twitches of the somatic musculature, including whisker twitches, and increased theta activity (Robinson et al., 1977; Montgomery et al., 2008). Paradoxically, despite the greater theta amplitude and frequency, we have previously shown that theta-gamma coupling is weaker in phasic than tonic REM sleep (Brankačk et al., 2012; Scheffzük et al., 2013). In light of the current results, we wondered whether the distinction in oscillatory LFP activity between phasic and tonic REM would relate to potential differences in breathing rate.

Figure 3A shows an example of a phasic REM sleep episode, characterized by a burst of increased theta amplitude and frequency. Phasic REM sleep comprised 2.48% ± 0.65% of REM sleep epochs (M ± SD over 22 mice; Figure 3B), a similar percentage to previous studies (Mizuseki et al., 2011; Brankačk et al., 2012; Scheffzük et al., 2013). In addition to increased theta activity, phasic REM also exhibited higher slow- and fast-gamma power than tonic REM sleep (Figure 3C), but reduced coupling between theta phase and the amplitude of both gamma sub-bands (Figure 3D,E; theta-fast gamma: t(21)=7.02, p<10^−6^; theta-slow gamma: t(21)=4.54, p<0.001, paired t-tests), thus consistent with previous studies (Brankačk et al., 2012; Scheffzük et al., 2013).

**Figure 3.**
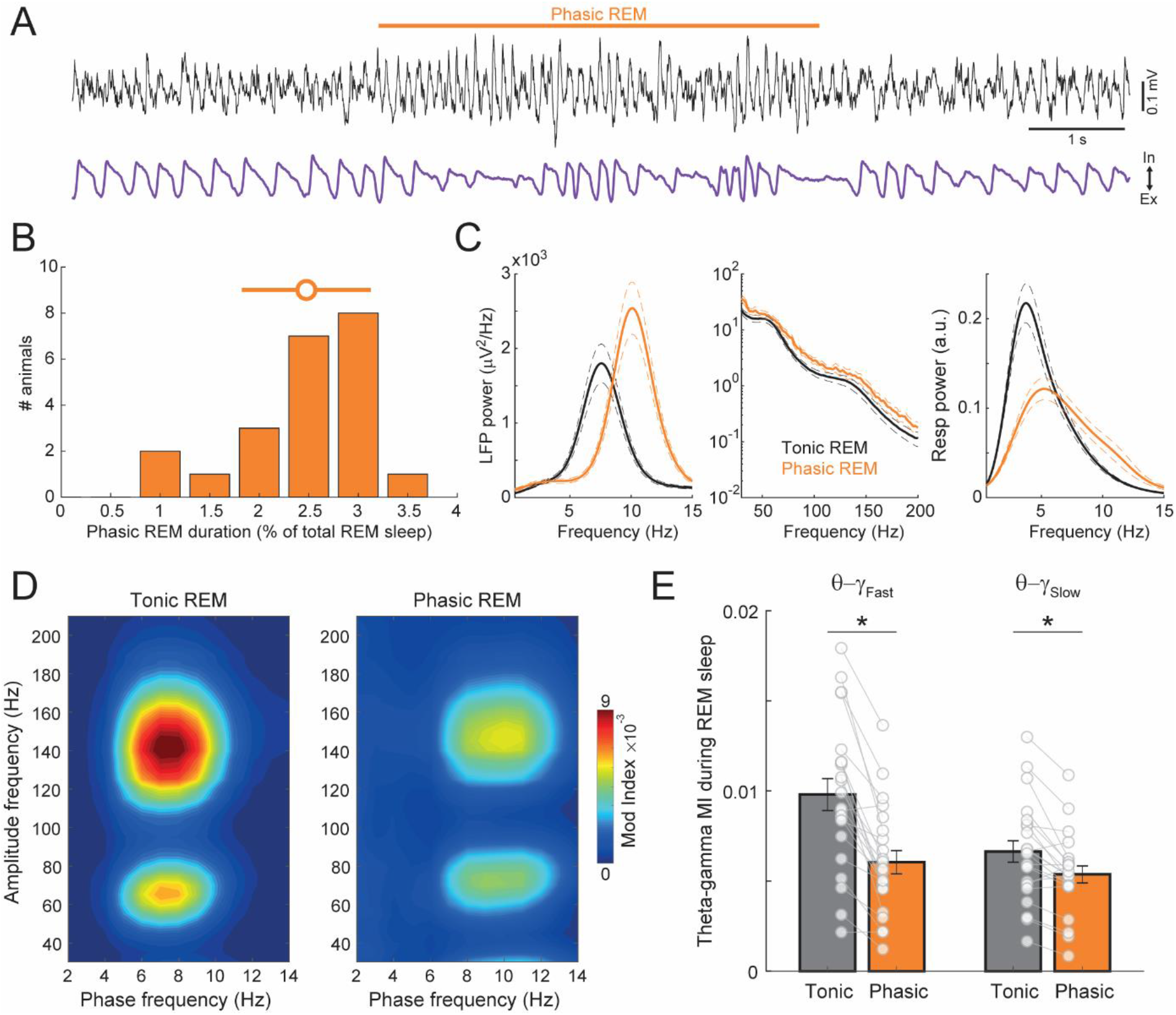
Phasic REM sleep has greater theta and gamma activity, but reduced theta-gamma coupling. (A) Example LFP and Resp traces during a phasic REM sleep episode amidst tonic REM sleep. (B) Histogram counts of the percentage of phase REM sleep per animal. (C) LFP (left and middle panels) and Resp (right) power spectra for tonic and phasic REM sleep (mean over 22 mice; dashed lines represent ± SEM). Notice higher theta and gamma power during phasic REM sleep. (D) Average comodulogram for tonic and phasic REM sleep epochs (n=22 mice). Notice faster gamma and theta frequencies and lower coupling strength during phasic REM. (E) Bar plots showing mean ± SEM theta-gamma coupling during tonic and phasic REM sleep. The connected circles show data for individual animals. *p<0.001, paired t-tests.

The power spectra of the plethysmograph-derived respiration signal in tonic and phasic REM sleep showed a large overlap, though phasic REM sleep exhibited a faster peak power frequency (tonic: 3.68 ± 0.29 Hz; phasic: 5.46 ± 1.43 Hz [M ± SD, n=22 mice]; t(21)=-5.77, p<10^−5^, paired t-test) and a fatter power tail towards theta and alpha frequencies (Figure 3C right). Accordingly, overlapping frequency distributions were also obtained in histogram counts of breathing rate over the pool of non-overlapping 500-ms windows (Figure 4A). The mean breathing rate over animals during tonic and phasic REM was 4.5 Hz ± 0.4 Hz and 6.1 Hz ± 0.92 Hz (M ± SD; n = 22 mice), respectively, a statistically significant difference (t(21)=10.14, p<10^−8^, paired t-test).

**Figure 4.**
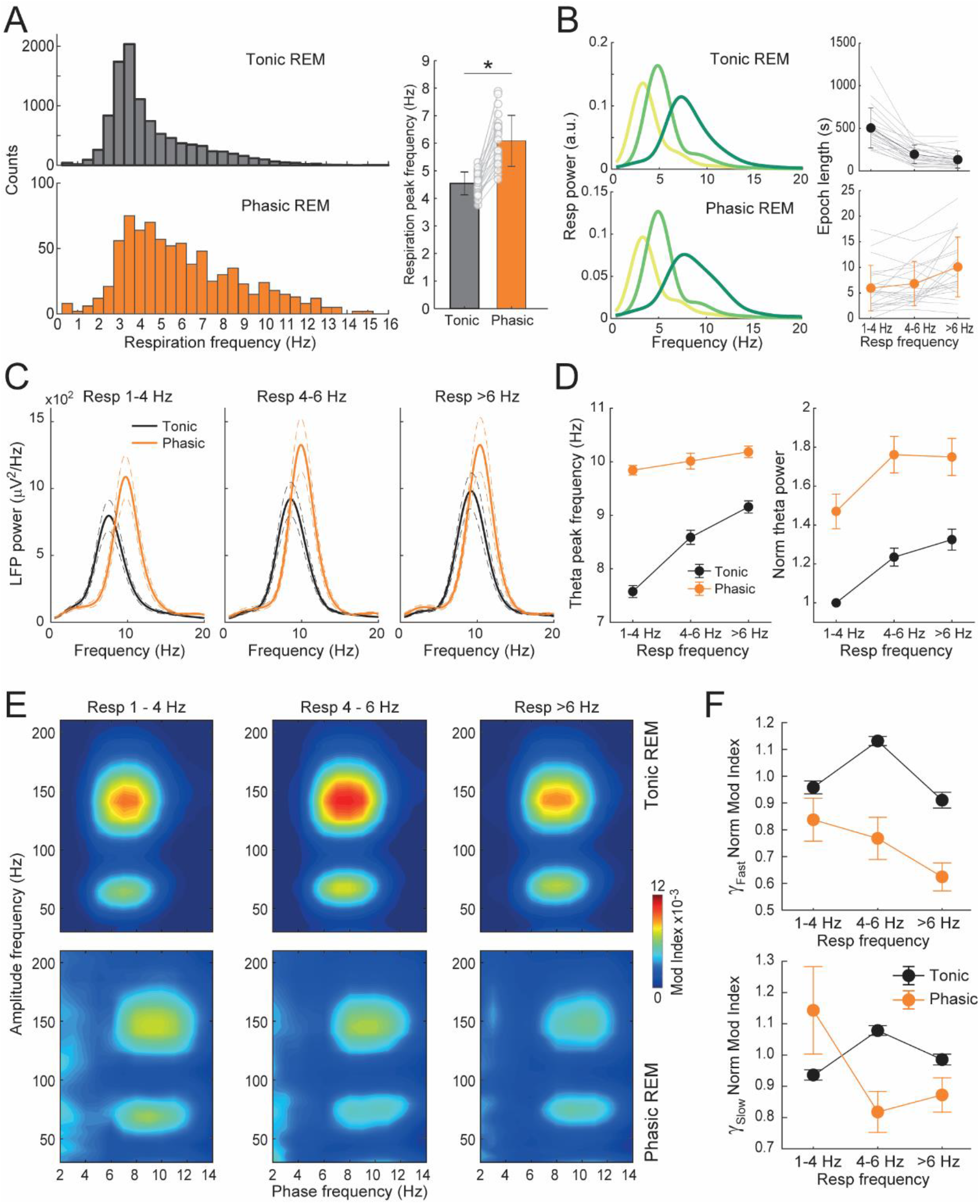
Breathing rate differentially modulates theta-gamma coupling in tonic and phasic REM sleep. (A) (Left) Histogram counts of respiration peak frequency in 500-ms windows within tonic (top) and phasic (bottom) REM sleep (pool over 22 animals). (Right) Mean (±SEM) breathing rate in each state; circles show individual animals. *p<10^−8^, paired t-test. (B) (Left) Mean Resp power spectra for respiration-binned epochs within tonic and phasic REM sleep (respiration frequency intervals: 1-4 Hz, 4-6 Hz, and >6 Hz). (Right) Total length of respiration-binned sub-epochs within each state (mean ±SEM; gray lines show individual animals). (C) Mean (±SEM) power spectra for the tonic and phasic REM sub-epochs binned by respiration frequency. (D) Theta frequency and normalized power as a function of breathing rate (mean ±SEM; n=22 mice). Power normalization was obtained by dividing by the theta power value of tonic REM in the first respiration frequency bin. (E) Average phase-amplitude comodulograms for the respiration frequency bins. Notice greater theta-gamma coupling during tonic REM sleep for all breathing rates. (F) Normalized theta-fast gamma (top) and theta-slow gamma (bottom) coupling strength for tonic and phasic REM as a function of breathing rate (mean ± SEM; n=22 mice). The normalization was performed within animals by dividing individual values by the mean modulation index during tonic REM. See text for statistical analyses.

To investigate the influence of breathing rate over oscillatory LFP activity in phasic and tonic REM sleep, we next adopted a similar approach as above to analyze LFP epochs restricted to certain respiration frequency intervals. However, given the sparsity of phasic REM episodes, to allow for a reliable analysis (i.e., long enough epochs) we employed fewer but wider respiration frequency bins: 1-4 Hz, 4-6 Hz and >6 Hz (Figure 4B). We found that theta power and frequency increased with increasing breathing rate within both tonic and phasic REM sleep (Figure 4C,D; theta power: F(2,120)=12.06, p<0.0001; theta frequency: F(2,120)=37.76, p<10^−12^, two-way ANOVA, column effect). Moreover, theta power and frequency were higher in phasic than tonic REM sleep irrespectively of breathing rate (Figure 4C,D; theta power: F(1,120)=69.49, p<10^−12^; theta frequency: F(1,120)=292.2, p<10^−33^, two-way ANOVA, row effect).

Interestingly, when computing phase-amplitude coupling binned by respiration frequency, we found that theta-fast gamma coupling was always higher for tonic than for phasic REM sleep irrespectively of breathing rate (Figure 4E,F; F(1,107)=47.45, p<10^−9^, two-way ANOVA, row effect). Furthermore, the inverted V-shaped relation (c.f. Figure 2C) was only found for tonic REM, while theta-fast gamma coupling strength during phasic REM linearly decreased with increasing breathing rate (Figure 4F top; F(2,107)=3.47, p=0.03, two-way ANOVA, interaction effect). A different profile of breathing rate dependence between phasic and tonic REM sleep was also observed for theta-slow gamma coupling, in which phasic REM had higher coupling strength for the slowest respiration range but otherwise lower values than tonic REM at faster breathing rates (Figure 4F bottom; F(2,107)=9.69, p=0.0001, two-way ANOVA, interaction effect).

In all, we conclude that the differences in theta-gamma coupling strength between phasic and tonic REM sleep are not explained by differences in breathing rate. Moreover, the results show that the breathing rate differently relates to oscillatory network activity in each state.

## Discussion

By focusing on studying phase-entrainment relations, a myriad of recent studies has recently demonstrated that nasal breathing is capable of driving LFP activity to oscillate phase-locked to respiration cycles in many non-olfactory regions (Tort et al., 2018a; Heck et al., 2019). They have also convincingly shown that theta and respiration tend not to have the same frequency and hence do not phase-lock (see Nguyen Chi et al., 2016 and Tort et al., 2018b for a discussion of breathing/sniffing at the same frequency as theta). Therefore, the growing consensus was that respiration and theta rhythms would be largely independent. Contrary to this idea, however, here we report an intriguing but very robust relation between respiration and theta (see supporting figures for individual results of each of the 22 animals). This relationship takes place not through phase-relations but between respiration rate and theta-gamma coupling strength.

The inverted V- or W-shaped dependence of theta-gamma coupling on breathing rate is admittedly unexpected, and in what follows we speculate on possible reasons for it. We first note that such an effect would not be due to any kind of oscillatory “resonance” phenomenon since the breathing frequency in which theta-gamma coupling is maximal is between 4 to 6 Hz, differing therefore from the theta peak frequency (7-9 Hz). Interestingly, fear conditioning studies have shown that mice tend to breathe around 4 Hz when exhibiting freezing (Bagur et al., 2018; Moberly et al., 2018), a behavior associated with fear memory retrieval. Moreover, another study in mice exposed to the plus maze found breathing rates of 2-4 Hz at basal levels, when animals rested still in the closed arms, but which reached 4-6 Hz when animals explored the open arms (Okonogi et al., 2018). It is thus possible that this particular breathing rate in mice marks an attentive brain state, which in this case would also be associated with highest rhythmic interactions of neocortical networks. Under this view, respiration rate and theta-gamma coupling would not necessarily be directly related but would both reflect an optimal level of network activity driven by a third player, most likely activating circuits in the brainstem. In fact, pontine nuclei have long been implicated in the onset and maintenance of REM sleep (Luppi et al., 2012; Peever and Fuller, 2017), and there is evidence for a direct communication between pontine cholinergic neurons and nearby circuits in brainstem nuclei that control respiration (Orem, 1980; Boutin et al., 2017; Benarroch, 2018; Del Negro et al., 2018).

Interestingly, even though the respiration rate exhibited a skewed distribution towards faster frequencies (Figure 1C), the mode frequency during sleep was ~3 Hz, with a typical, “baseline” range between 2 and 4 Hz (Figures 2A and 4A top left). This means that the breathing rates between 4 and 6 Hz associated with the highest theta-gamma coupling were much less common (see Figure 4B top right), and as such can be considered deviations from the basal respiratory activity during sleep. Typically, changes in network oscillations have been related to particular behaviors, but here, the animals were in the behaviorally passive state of REM sleep throughout the experiment. Nevertheless, previous studies have shown that the micro-architecture of REM sleep is not uniform but rather exhibits different patterns of activity (Bandarabadi et al., 2019; Blumberg et al., 2020; Simor et al., 2020). These, in turn, would be reflected in different sub-patterns of network activity when examined at a more detailed scale as performed here.

Understanding the biological functions of REM sleep has been an active research field for decades (Siegel, 2005). Among the proposed functions, REM sleep has been linked to regulation of brain temperature (Wehr, 1992), synaptic plasticity (Dumoulin Bridi et al., 2015; Li et al., 2017), learning and memory consolidation (Rasch and Born, 2013; Boyce et al., 2016), and emotional regulation (Walker, 2010; Goldstein and Walker, 2014; Tempesta et al., 2018). Regarding the latter, it is interesting to note that respiration patterns also change with emotional state (Grossman and Wientjes, 2001; Philippot et al., 2002; Zhang et al., 2017), and, in fact, a subpopulation of glutamatergic neurons in the preBötzinger complex has been shown to influence arousal and emotional regulation (Yackle et al., 2017; Del Negro et al., 2018). It is often assumed that rodents experience dreams similar to humans, although this is difficult to prove (e.g., Louie and Wilson, 2001; see also Manger and Siegel, 2020). In any case, it is well feasible that breathing rate changes during REM sleep relate to the animals experiencing different emotional contents. In this sense, an arousal-like sensation leading to mildly faster breathing frequencies could set the network to an optimal functional state, while large increases of breathing rate would associate with a strong emotional state and be detrimental for proper information processing.

Previous studies have shown that theta-gamma coupling is strongest during REM sleep (Scheffzük et al., 2011; Brankačk et al., 2012; Schomburg et al., 2014; Koike et al., 2017; Scheffer-Teixeira and Tort, 2017; Bandarabadi et al., 2019). This is especially the case of neocortical theta-fast gamma coupling, which oftentimes is not detected during awake behaviors and is thus believed to play functional roles in sleep-dependent information processing (Scheffzük et al., 2011; Tort et al., 2013; Bandarabadi et al., 2019). Interestingly, a recent study has called attention to the fact that the strength of theta-gamma coupling during REM sleep is not constant but varies over time, although the possible factors accounting for this fluctuation were not identified (Bandarabadi et al., 2019). The present results shed light on at least one such factor, namely the breathing rate. Nevertheless, as discussed above, at present it is unclear whether changes in breathing rate would directly relate to theta-gamma coupling or rather be driven by changes in arousal levels that would, in turn, affect both rhythmic breathing and network activity. In addition, as shown by previous research (Brankačk et al., 2012; Scheffzük et al., 2013) and here (Figure 3), another important factor accounting for differences in theta-gamma coupling strength is whether the animal is displaying tonic or phasic REM sleep, which we discuss next.

REM sleep can be subdivided into tonic and phasic states on the basis of behavioral and electrophysiological variables (see Simor et al., 2020 for a recent review). The tonic state comprises most of REM sleep epochs; it is considered a more quiescent state that can be briefly interrupted by bursts of phasic phenomena that occur many times along REM sleep (Robinson et al., 1977; Simor et al., 2020). The phasic state corresponds to the short periods where eye movements actually occur, and which further associates with ponto-geniculo-occipital waves, clonic twitches of the skeletal musculature, as well as with changes in cardiac and respiratory activity (Robinson et al., 1977; Simor et al., 2020). In rodents, both REM sub-states occur along with theta oscillations (Robinson et al., 1977; Montgomery et al., 2008), but, interestingly, theta activity during tonic REM sleep depends on acetylcholine (atropine sensitive) while theta during phasic REM does not (atropine resistant) (Robinson et al., 1977). Tonic REM sleep is sometimes compared to a more “relaxed” wakeful-like brain state, while phasic REM sleep would correspond to a more activated state (De Carli et al., 2016), though the brain is much less susceptible to perturbations by sensory stimuli during phasic REM (Price and Kremen, 1980; Wehrle et al., 2007; Simor et al., 2020).

We have previously shown that even though neocortical LFP activity during phasic REM sleep exhibits higher theta and gamma power compared to tonic REM sleep, somewhat paradoxically, this state is characterized by lower theta-gamma coupling (Brankačk et al., 2012; Scheffzük et al., 2013), a result we were able to corroborate here (Figure 3D,E). Curiously, in the seminal report describing REM sleep in humans (Aserinsky and Kleitman, 1953), the authors noticed that breathing rate was faster during ocular movements (i.e., during phasic REM sleep), thus consistent with the current findings in mice (Figure 4A). In light of the relationship between breathing rate and theta-gamma coupling described here (Figure 2), a key question was to assess whether the differences in theta-gamma coupling strength between tonic and phasic REM sleep could be accounted for by differences in breathing rate. Even though animals tend to breathe faster during phasic REM, our results show that this does not explain the lower coupling strength. Rather, we found that breathing rate influences oscillatory network activity in each sub-state, but in specific manners, and that differences between phasic and tonic REM sleep also occur within fixed respiration frequency ranges (Figure 4). Interestingly, they further show that breathing differentially relates to theta-gamma coupling within each REM sleep state (Figure 4E,F). For instance, the inverted V-shaped relation for theta-fast gamma coupling and breathing rate only takes place during tonic REM sleep, while for phasic REM theta-fast gamma coupling linearly decreases with faster breathing (Figure 4F top). (Notice that the inverted V-shaped relation appears when analyzing all REM sleep epochs due to the abundance of the tonic state).

In summary, our results reveal that the respiratory activity correlates with brain activity not only through the well-described phase-entrainment effects, but also through an intriguing relation involving the instantaneous breathing rate and theta-gamma coupling. In addition, our results further support physiological and functional distinctions between tonic and phasic REM sleep states.

## Materials and Methods

### Ethics statement

This study was approved by the Governmental Supervisory Panel on Animal Experiments of Baden-Württemberg (35-9185.81 G84/13 and G-115/14). The present experiments were carried out in line with the guidelines of the European Science Foundation (2000) and the US National Institute of Health Guide for the Care and Use of Laboratory Animals (2011).

### Animal care

A total of twenty-two C57/Bl6 mice were used, nine females and thirteen males (body weight: 22-37 g; age: 13-19 weeks). The animals were housed in groups of 4 with access to food and water *ad libitum*. After electrode implantation, animals were kept individually until completion of the experiments. Mice were housed on a 12-hour light-dark-cycle (lights off at 8:00 a.m., except 2 animals on the opposite circadian phase).

### Surgery

Mice were implanted with epidural surface electrodes. Before surgery, buprenorphine (0.1 mg/kg) was administered for pain control. This was repeated after completion of the surgery and every 8 h if necessary. Surgery was performed under isoflurane anesthesia (4% for induction, 1.5% during surgery; for further details, see Jessberger et al., 2016 and Zhang et al., 2016). After exposing the skull, 0.5 to 1 mm diameter holes were drilled above the parietal cortex according to stereotactic coordinates (2 mm posterior bregma, 1.5 mm lateral to the midline). Reference and ground electrodes were screwed into the skull above the cerebellum. During surgery, body temperature was monitored and maintained at 37-38 °C. Electrodes consisted of stainless steel watch screws. Animals were allowed 7 days of recovery prior to experiments.

### Electrophysiological recordings and behavioral staging

Mice were placed in a whole-body plethysmograph (EMKA Technologies, SAS, France), which was customized to allow simultaneous recordings of local field potentials (LFPs). It consisted of a transparent cylinder (78 mm inner diameter, 165 mm height) connected to a reference chamber (Figure 1A). Individual recording sessions lasted from 2 to 6 hours (mean: 3 h). Monopolar electrophysiological signals were filtered (1-500Hz), amplified and digitized with a sampling frequency of 2.5 kHz (RHA2000, Intan Technologies, Los Angeles, USA). A three-dimensional accelerometer was custom-mounted on the amplifier board located on the head of the mice to allow movement detection. The three signals of the accelerometer were fed to three channels of the amplifier using the same band-pass filter as for the LFPs, therefore removing the gravity-induced sustained potentials of the accelerometers.

REM-sleep was manually staged by a senior researcher in the field (J.B.) and identified as follows: 1) minimal accelerometer activity and 2) continuous theta rhythm in the parietal cortex following a sleep stage with slow-waves typical for non-REM sleep (Brankačk et al., 2010).

### Data analysis

We used built-in and custom-written routines in MatLab^©^ (The Mathworks Inc., Natick, MA). For each animal, all identified REM sleep epochs were first detrended and then concatenated before analysis.

### Spectral analysis and estimation of breathing rate

Standard, Fourier-based power spectra were computed using the Welch periodogram through the pwelch.m function from the Signal Processing Toolbox. In Figure 1C, we used 4-s windows with 50% overlap. Time-frequency power spectrograms of respiratory activity (derived from the plethysmograph signal) were computed employing the spectrogram.m function from the Signal Processing Toolbox, using 500-ms windows with no overlap. The numerical frequency resolution was set to 0.05 Hz. For each 500-ms window, the “instantaneous” breathing rate was taken as the frequency with maximal power.

### Cross-frequency coupling

Phase-amplitude comodulograms were computed using the Modulation Index (MI) as described in Scheffer-Teixeira and Tort (2018) and available at https://github.com/tortlab/phase-amplitude-coupling. Filtering was obtained by means of the eegfilt.m function from the EEGLAB toolbox (Delorme and Makeig, 2004; http://sccn.ucsd.edu/eeglab/), which relies on a linear finite impulse response filter applied in the forward and backward directions to nullify phase delays. After filtering, the phase and amplitude time series were obtained from the analytical signal representation based on the Hilbert transform (hilbert.m function). Filtering bandwidths were 4 Hz for phase frequencies (1 Hz step; center frequencies from 2 to 14 Hz) and 20 Hz for amplitude frequencies (5 Hz step; center frequencies from 30 to 210 Hz). The phase time series were binned into eighteen 20° non-overlapping phase bins. In Figure 1E, we computed phase-amplitude comodulograms using all REM sleep epochs. For each frequency pair of interest (i.e., theta-slow gamma or theta-fast gamma), coupling strength was taken from the comodulogram as the maximal MI value in the corresponding frequency ROI (i.e., 4-12 Hz vs 50-80 Hz or 4-12 Hz vs 120-160 Hz). For computing the “respiration-binned” comodulograms (Figures 2B and 4E), for each analyzed respiration frequency interval we first identified all 500-ms periods with breathing rate within that interval (as assessed by the peak power of the respiration signal in 500-ms non-overlapping windows; see subsection above). The corresponding comodulogram was then computed using phase and amplitude data restricted to the identified periods. Of note, to avoid edge effects, we first filtered the LFP and obtained the amplitude and phase time series, and only afterward we extracted the periods of interest for MI computation (Tort et al., 2008). As before, theta-slow gamma and theta-fast gamma coupling strength were taken as the maximal MI value in the corresponding frequency ROI of the comodulogram.

Since phase-amplitude coupling estimation has a positive bias for very short epochs (Tort et al., 2010), for each respiration frequency interval in Figures 2 we only considered animals whose total analyzed length was >10 s. In practice, the sample size was all the 22 mice for all respiration intervals except for two bins: resp between 9 and 11 Hz (n=20 mice) and resp >10 Hz (n= 17 mice).

### Detection of phasic REM sleep

Bursts of phasic REM sleep were detected as in previous work (Mizuseki et al., 2011; Brankačk et al., 2012): first, the REM sleep LFP was band-pass filtered between 4 and 12 Hz to obtain the timestamps of the theta peaks. Interpeak intervals were smoothed using an 11-sample moving average. Candidate epochs were those in which the smoothed interpeak intervals were in the 10^th^ percentile. Next, the following criteria applied: 1) minimal duration of 900 ms; 2) minimal smoothed interpeak interval during the candidate epoch shorter than the 5^th^ percentile; and 3) mean theta amplitude during the candidate epoch larger than the mean theta amplitude across all REM sleep. Following this definition, phasic REM corresponded to an average of 2.48% of all REM sleep (see Figure 3B), a similar percentage as in our previous study (Brankačk et al., 2012).

For the purpose of the analyses performed in Figures 3 and 4, tonic REM sleep was considered as the remaining REM sleep after excluding all phasic REM sleep periods as well as 2-s before and after them.

In Figure 3C, power spectra for tonic and phasic REM sleep were obtained using 0.5-s windows and 50% overlap. The comodulograms in Figures 3D and 4E were computed using the REM sleep epochs restricted to the corresponding sub-state (phasic or tonic). In Figure 4F, for each respiration interval we only considered animals whose total analyzed length was >4 s; this resulted in n=22 mice for all intervals in the tonic group, and n=13, 15 and 19 mice in the phasic group for the 1^st^, 2^nd^ and 3^rd^ respiration interval, respectively.

### Statistical analysis

Data are expressed as mean ± SEM. Electrophysiological measures were compared using either paired t-tests, repeated measures ANOVA or two-way ANOVA, as appropriate and stated in the main text or figure legends. Statistical significance was set at alpha = 0.05.

## Acknowledgments

This work was supported by the Deutsche Forschungsgemeinschaft (SFB 1134/A01; Dr 326/10-1), Bundesministerium für Bildung und Forschung (German-Brazil Cooperation grant No. 01DN12098), the Brazilian National Council for Scientific and Technological Development (CNPq), the Brazilian Coordination for the Improvement of Higher Education Personal (CAPES), and the Alexander von Humboldt Foundation. The funders had no role in study design, data collection and analysis, decision to publish, or preparation of the manuscript.

## Supplementary Material

**Figure S1.**
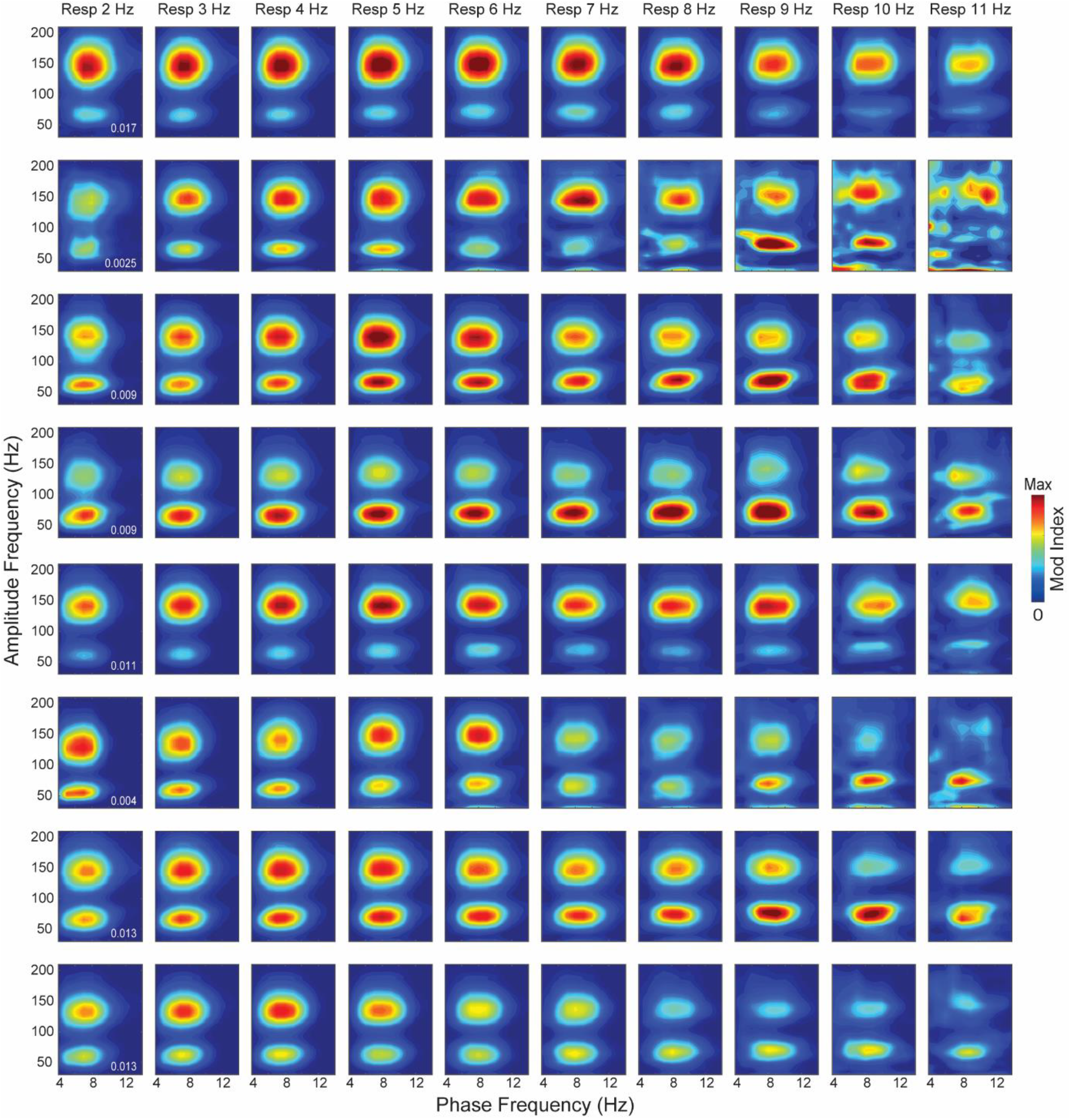
Comodulation plots for individual animals (1/3). Phase-amplitude comodulograms computed for REM sleep LFP epochs binned by respiration frequency. The title of each column states the average breathing rate (respiration frequency intervals: 1-3 Hz, 2-4 Hz, 3-5 Hz, 4-6 Hz, 5-7 Hz, 6-8 Hz, 7-9 Hz, 8-10 Hz, 9-11 Hz, and >10 Hz). Each row shows the results of a different mouse (animals #1 to #8 shown in this figure). The inset numbers in the leftmost comodulograms indicate the maximum of the modulation index color scale for the corresponding row. Notice dependence of theta-gamma coupling strength on breathing rate for all animals.

**Figure S2.**
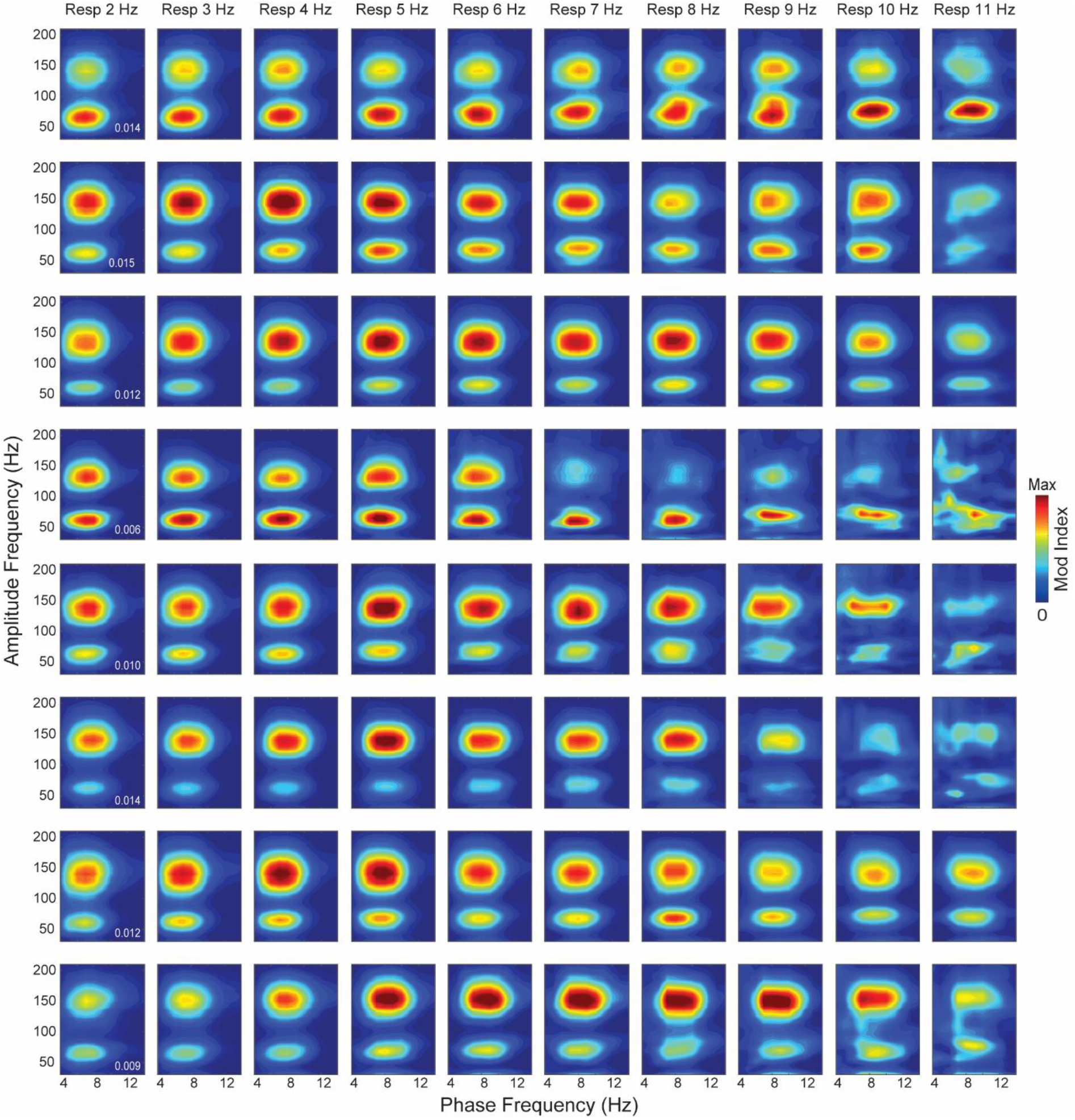
Comodulation plots for individual animals (2/3). As in Figure S1, but for animals #9 to #16.

**Figure S3.**
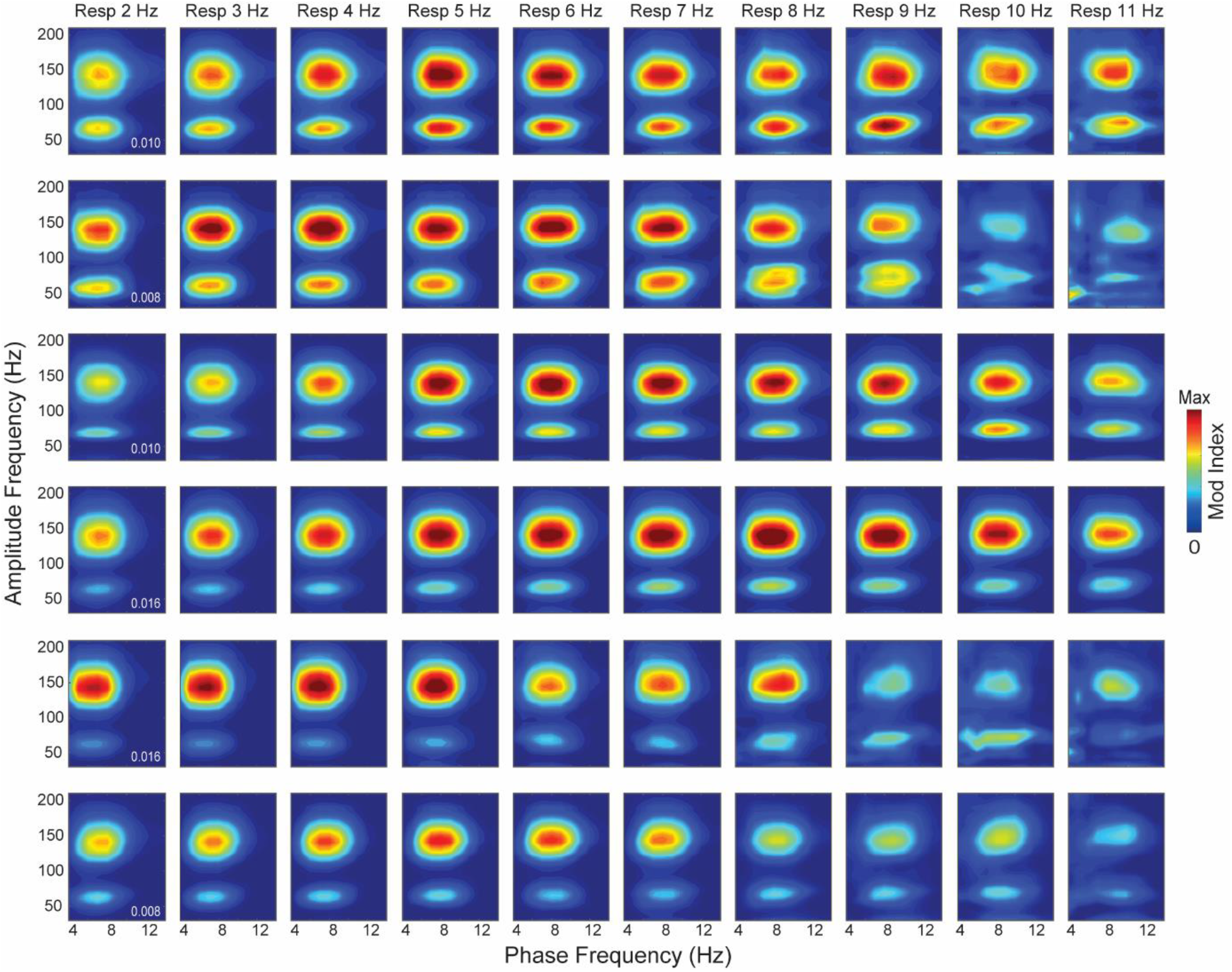
Comodulation plots for individual animals (3/3). As in Figure S1, but for animals #17 to #22.

## References

Aserinsky E, Kleitman N (1953) Regularly occurring periods of eye motility, and concomitant phenomena, during sleep. Science 118:273–274.

Bagur S, Lefort J, Lacroix M, de Lavilléon G, Herry C, Billand C, Geoffroy H, Benchenane K (2018) Dissociation of fear initiation and maintenance by breathing-driven prefrontal oscillations. bioRxiv.

Bandarabadi M, Boyce R, Gutierrez Herrera C, Bassetti CL, Williams S, Schindler K, Adamantidis A (2019) Dynamic modulation of theta-gamma coupling during rapid eye movement sleep. Sleep 42:zsz182.

Belluscio MA, Mizuseki K, Schmidt R, Kempter R, Buzsáki G (2012) Cross-frequency phase-phase coupling between θ and γ oscillations in the hippocampus. J Neurosci 32:423–435.

Benarroch EE (2018) Brainstem integration of arousal, sleep, cardiovascular, and respiratory control. Neurology 91:958–966.

Biskamp J, Bartos M, Sauer J-F (2017) Organization of prefrontal network activity by respiration-related oscillations. Sci Rep 7:45508.

Blumberg MS, Lesku JA, Libourel P-A, Schmidt MH, Rattenborg NC (2020) What Is REM Sleep? Curr Biol 30:R38–R49.

Bott J-B, Muller M-A, Jackson J, Aubert J, Cassel J-C, Mathis C, Goutagny R (2016) Spatial reference memory is associated with modulation of theta-gamma coupling in the dentate gyrus. Cereb Cortex 26:3744–3753.

Boutin RCT, Alsahafi Z, Pagliardini S (2017) Cholinergic modulation of the parafacial respiratory group. J Physiol 595:1377–1392.

Boyce R, Glasgow SD, Williams S, Adamantidis A (2016) Causal evidence for the role of REM sleep theta rhythm in contextual memory consolidation. Science 352:812–816.

Brankačk J, Kukushka VI, Vyssotski AL, Draguhn A (2010) EEG gamma frequency and sleep-wake scoring in mice: comparing two types of supervised classifiers. Brain Res 1322:59–71.

Brankačk J, Scheffzük C, Kukushka VI, Vyssotski AL, Tort ABL, Draguhn A (2012) Distinct features of fast oscillations in phasic and tonic rapid eye movement sleep. J Sleep Res 21:630–633.

Buzsáki G, Wang X-J (2012) Mechanisms of Gamma Oscillations. Annu Rev Neurosci 35:203–225.

Canolty RT, Knight RT (2010) The functional role of cross-frequency coupling. Trends Cogn Sci 14:506–515.

Cavelli M, Castro-Zaballa S, Gonzalez J, Rojas-Líbano D, Rubido N, Velásquez N, Torterolo P (2020) Nasal respiration entrains neocortical long-range gamma coherence during wakefulness. Eur J Neurosci 51:1463–1477.

Chrobak JJ, Buzsáki G (1998) Gamma oscillations in the entorhinal cortex of the freely behaving rat. J Neurosci 18:388–398.

Colgin LL (2011) Oscillations and hippocampal-prefrontal synchrony. Curr Opin Neurobiol 21:467–474.

De Carli F, Proserpio P, Morrone E, Sartori I, Ferrara M, Gibbs SA, De Gennaro L, Lo Russo G, Nobili L (2016) Activation of the motor cortex during phasic rapid eye movement sleep. Ann Neurol 79:326–330.

Del Negro CA, Funk GD, Feldman JL (2018) Breathing matters. Nat Rev Neurosci 19:351–367.

Delorme A, Makeig S (2004) EEGLAB: an open source toolbox for analysis of single-trial EEG dynamics including independent component analysis. J Neurosci Methods 134:9–21.

Dumoulin Bridi MC, Aton SJ, Seibt J, Renouard L, Coleman T, Frank MG (2015) Rapid eye movement sleep promotes cortical plasticity in the developing brain. Sci Adv 1:e1500105.

Goldstein AN, Walker MP (2014) The role of sleep in emotional brain function. Annu Rev Clin Psychol 10:679–708.

Grossman P, Wientjes CJ (2001) How breathing adjusts to mental and physical demands. In: Respiration and emotion, pp 43–54. Springer.

Heck DH, Kozma R, Kay LM (2019) The rhythm of memory: how breathing shapes memory function. J Neurophysiol 122:563–571.

Heck DH, McAfee SS, Liu Y, Babajani-Feremi A, Rezaie R, Freeman WJ, Wheless JW, Papanicolaou AC, Ruszinkó M, Sokolov Y, Kozma R (2017) Breathing as a fundamental rhythm of brain function. Front Neural Circuits 10:115.

High Level Expert Group on Biology and Society, European Science Foundation, Strasbourg, France (2000) European Science Foundation policy briefing: use of animals in research. Altern Lab Anim ATLA 28:743–749.

Hyafil A, Giraud A-L, Fontolan L, Gutkin B (2015) Neural cross-frequency coupling: connecting architectures, mechanisms, and functions. Trends Neurosci 38:725–740.

Isomura Y, Sirota A, Özen S, Montgomery S, Mizuseki K, Henze DA, Buzsáki G (2006) Integration and Segregation of Activity in Entorhinal-Hippocampal Subregions by Neocortical Slow Oscillations. Neuron 52:871–882.

Ito J, Roy S, Liu Y, Cao Y, Fletcher M, Lu L, Boughter JD, Grün S, Heck DH (2014) Whisker barrel cortex delta oscillations and gamma power in the awake mouse are linked to respiration. Nat Commun 5:3572.

Jessberger J, Zhong W, Brankačk J, Draguhn A (2016) Olfactory Bulb Field Potentials and Respiration in Sleep-Wake States of Mice. Neural Plast 2016:4570831.

Koike BDV, Farias KS, Billwiller F, Almeida-Filho D, Libourel P-A, Tiran-Cappello A, Parmentier R, Blanco W, Ribeiro S, Luppi P-H, Queiroz CM (2017) Electrophysiological Evidence That the Retrosplenial Cortex Displays a Strong and Specific Activation Phased with Hippocampal Theta during Paradoxical (REM) Sleep. J Neurosci 37:8003–8013.

Lasztóczi B, Klausberger T (2014) Layer-specific GABAergic control of distinct gamma oscillations in the CA1 hippocampus. Neuron 81:1126–1139.

Li W, Ma L, Yang G, Gan W-B (2017) REM sleep selectively prunes and maintains new synapses in development and learning. Nat Neurosci 20:427–437.

Lisman JE, Jensen O (2013) The θ-γ neural code. Neuron 77:1002–1016.

Lockmann ALV, Laplagne DA, Leão RN, Tort ABL (2016) A respiration-coupled rhythm in the rat hippocampus independent of theta and slow oscillations. J Neurosci 36:5338–5352.

Lopes-Dos-Santos V, van de Ven GM, Morley A, Trouche S, Campo-Urriza N, Dupret D (2018) Parsing Hippocampal Theta Oscillations by Nested Spectral Components during Spatial Exploration and Memory-Guided Behavior. Neuron 100:940–952.e7.

Louie K, Wilson MA (2001) Temporally Structured Replay of Awake Hippocampal Ensemble Activity during Rapid Eye Movement Sleep. Neuron 29:145–156.

Luppi P-H, Clement O, Sapin E, Peyron C, Gervasoni D, Léger L, Fort P (2012) Brainstem mechanisms of paradoxical (REM) sleep generation. Pflugers Arch 463:43–52.

Manger PR, Siegel JM (2020) Do all mammals dream? J Comp Neurol 528:3198–3204.

Mizuseki K, Diba K, Pastalkova E, Buzsáki G (2011) Hippocampal CA1 pyramidal cells form functionally distinct sublayers. Nat Neurosci 14:1174–1181.

Moberly AH, Schreck M, Bhattarai JP, Zweifel LS, Luo W, Ma M (2018) Olfactory inputs modulate respiration-related rhythmic activity in the prefrontal cortex and freezing behavior. Nat Commun 9:1528.

Montgomery SM, Sirota A, Buzsáki G (2008) Theta and gamma coordination of hippocampal networks during waking and rapid eye movement sleep. J Neurosci 28:6731–6741.

National Research Council (US) Committee for the Update of the Guide for the Care and Use of Laboratory Animals (2011) Guide for the Care and Use of Laboratory Animals, 8th ed. Washington (DC): National Academies Press (US).

Nguyen Chi V, Müller C, Wolfenstetter T, Yanovsky Y, Draguhn A, Tort ABL, Brankačk J (2016) Hippocampal respiration-driven rhythm distinct from theta oscillations in awake mice. J Neurosci 36:162–177.

Okonogi T, Nakayama R, Sasaki T, Ikegaya Y (2018) Characterization of Peripheral Activity States and Cortical Local Field Potentials of Mice in an Elevated Plus Maze Test. Front Behav Neurosci 12:62.

Orem J (1980) Medullary respiratory neuron activity: relationship to tonic and phasic REM sleep. J Appl Physiol 48:54–65.

Peever J, Fuller PM (2017) The Biology of REM Sleep. Curr Biol CB 27:R1237–R1248.

Philippot P, Chapelle G, Blairy S (2002) Respiratory feedback in the generation of emotion. Cogn Emot 16:605–627.

Price LJ, Kremen I (1980) Variations in behavioral response threshold within the REM period of human sleep. Psychophysiology 17:133–140.

Rasch B, Born J (2013) About sleep’s role in memory. Physiol Rev 93:681–766.

Robinson TE, Kramis RC, Vanderwolf CH (1977) Two types of cerebral activation during active sleep: relations to behavior. Brain Res 124:544–549.

Rojas-Líbano D, Wimmer Del Solar J, Aguilar-Rivera M, Montefusco-Siegmund R, Maldonado PE (2018) Local cortical activity of distant brain areas can phase-lock to the olfactory bulb’s respiratory rhythm in the freely behaving rat. J Neurophysiol 120:960–972.

Sakai K, Sano K, Iwahara S (1973) Eye movements and hippocampal theta activity in cats. Electroencephalogr Clin Neurophysiol 34:547–549.

Scheffer-Teixeira R, Belchior H, Caixeta FV, Souza BC, Ribeiro S, Tort ABL (2012) Theta phase modulates multiple layer-specific oscillations in the CA1 region. Cereb Cortex 22:2404–2414.

Scheffer-Teixeira R, Tort AB (2018) Theta-Gamma Cross-Frequency Analyses (Hippocampus). Encycl Comput Neurosci:1–15.

Scheffer-Teixeira R, Tort ABL (2017) Unveiling fast field oscillations through comodulation. eNeuro 4:ENEURO. 0079-17.2017.

Scheffzük C, Kukushka VI, Vyssotski AL, Draguhn A, Tort ABL, Brankačk J (2011) Selective coupling between theta phase and neocortical fast gamma oscillations during REM-sleep in mice. PloS One 6:e28489.

Scheffzük C, Kukushka VI, Vyssotski AL, Draguhn A, Tort ABL, Brankačk J (2013) Global slowing of network oscillations in mouse neocortex by diazepam. Neuropharmacology 65:123–133.

Schomburg EW, Fernández-Ruiz A, Mizuseki K, Berényi A, Anastassiou CA, Koch C, Buzsáki G (2014) Theta phase segregation of input-specific gamma patterns in entorhinal-hippocampal networks. Neuron 84:470–485.

Siegel JM (2005) Clues to the functions of mammalian sleep. Nature 437:1264–1271.

Simor P, van der Wijk G, Nobili L, Peigneux P (2020) The microstructure of REM sleep: Why phasic and tonic? Sleep Med Rev 52:101305.

Sirota A, Montgomery S, Fujisawa S, Isomura Y, Zugaro M, Buzsáki G (2008) Entrainment of neocortical neurons and gamma oscillations by the hippocampal theta rhythm. Neuron 60:683–697.

Stujenske JM, Likhtik E, Topiwala MA, Gordon JA (2014) Fear and safety engage competing patterns of theta-gamma coupling in the basolateral amygdala. Neuron 83:919–933.

Tempesta D, Socci V, De Gennaro L, Ferrara M (2018) Sleep and emotional processing. Sleep Med Rev 40:183–195.

Tort ABL, Brankačk J, Draguhn A (2018a) Respiration-Entrained Brain Rhythms Are Global but Often Overlooked. Trends Neurosci 41:186–197.

Tort ABL, Komorowski R, Eichenbaum H, Kopell N (2010) Measuring phase-amplitude coupling between neuronal oscillations of different frequencies. J Neurophysiol 104:1195–1210.

Tort ABL, Komorowski RW, Manns JR, Kopell NJ, Eichenbaum H (2009) Theta-gamma coupling increases during the learning of item-context associations. Proc Natl Acad Sci U S A 106:20942–20947.

Tort ABL, Kramer MA, Thorn C, Gibson DJ, Kubota Y, Graybiel AM, Kopell NJ (2008) Dynamic cross-frequency couplings of local field potential oscillations in rat striatum and hippocampus during performance of a T-maze task. Proc Natl Acad Sci U S A 105:20517–20522.

Tort ABL, Ponsel S, Jessberger J, Yanovsky Y, Brankačk J, Draguhn A (2018b) Parallel detection of theta and respiration-coupled oscillations throughout the mouse brain. Sci Rep 8:6432.

Tort ABL, Rotstein HG, Dugladze T, Gloveli T, Kopell NJ (2007) On the formation of gamma-coherent cell assemblies by oriens lacunosum-moleculare interneurons in the hippocampus. Proc Natl Acad Sci U S A 104:13490–13495.

Tort ABL, Scheffer-Teixeira R, Souza BC, Draguhn A, Brankačk J (2013) Theta-associated high-frequency oscillations (110-160Hz) in the hippocampus and neocortex. Prog Neurobiol 100:1–14.

Walker MP (2010) Sleep, memory and emotion. Prog Brain Res 185:49–68.

Wehr TA (1992) A brain-warming function for REM sleep. Neurosci Biobehav Rev 16:379–397.

Wehrle R, Kaufmann C, Wetter TC, Holsboer F, Auer DP, Pollmächer T, Czisch M (2007) Functional microstates within human REM sleep: first evidence from fMRI of a thalamocortical network specific for phasic REM periods. Eur J Neurosci 25:863–871.

Yackle K, Schwarz LA, Kam K, Sorokin JM, Huguenard JR, Feldman JL, Luo L, Krasnow MA (2017) Breathing control center neurons that promote arousal in mice. Science 355:1411–1415.

Yanovsky Y, Ciatipis M, Draguhn A, Tort ABL, Brankačk J (2014) Slow oscillations in the mouse hippocampus entrained by nasal respiration. J Neurosci 34:5949–5964.

Zhang L, Lee J, Rozell C, Singer AC (2019) Sub-second dynamics of theta-gamma coupling in hippocampal CA1. eLife 8:e44320.

Zhang Q, Chen X, Zhan Q, Yang T, Xia S (2017) Respiration-based emotion recognition with deep learning. Comput Ind 92–93:84–90.

Zhang X, Zhong W, Brankačk J, Weyer SW, Müller UC, Tort ABL, Draguhn A (2016) Impaired theta-gamma coupling in APP-deficient mice. Sci Rep 6:21948.

Zhong W, Ciatipis M, Wolfenstetter T, Jessberger J, Müller C, Ponsel S, Yanovsky Y, Brankačk J, Tort ABL, Draguhn A (2017) Selective entrainment of gamma subbands by different slow network oscillations. Proc Natl Acad Sci U S A 114:4519–4524.

